# Population colonization history influences behavioral responses of European starlings in personality tests

**DOI:** 10.1101/2021.07.24.453662

**Authors:** Alexandra Rodriguez, Martine Hausberger, Patricia Le Quilliec, Philippe Clergeau, Laurence Henry

## Abstract

To understand the processes involved in biological invasions, the genetic, morphological, physiological and behavioral characteristics of invasive populations need to be understood.

Many invasive species have been reported to be flying species. In birds, both invaders and migrants encounter novel situations, therefore one could expect that both groups might react similarly to novelty.

Here we analyzed the behavioral responses of individuals from three populations of European starling *Sturnus vulgaris*: a population settled for centuries in a rural region, a population that recently colonized an urban area, and a population of winter migrant birds. We conducted a social isolation test, a novel environment test, a novel food test and a novel object test to explore their reactions towards novelty. We identified and characterized different behavioral profiles for each test.

The group of migratory adults appeared to be less anxious in social isolation than the group of urban young. Urban and migrant groups entered the novel environment sooner than rural birds. Shy, bold and intermediate individuals were observed in all three groups when presented with novel food. Finally, the proportion of shy individuals which did not touch the novel object was higher than the proportion of bold individuals in the rural group. Our study emphasizes that neophilia or boldness present in migrant and invasive populations may facilitate the occupation of novel habitats. Our analysis also suggests that mixed reactions of neophobia ensure behavioral flexibility in a gregarious invasive species.

**Significant statement:** In this paper, we show that an invasive species like European starling, *Sturnus vulgaris*, presents an important flexibility in neophobia and in reaction towards social isolation. These variations depend on the settlement history of populationseven when the birds had been wild-caught as nestlings and hand-raised in standard conditions. This is significant because it highlights possible scenarios of colonization processes.

We believe that this manuscript is appropriate for publication by Behavioral Ecology and Sociobiology because it places individuals’ behavior in the core mechanisms of an ecological phenomenom as biological invasions. Our manuscript enlarges the paradigms related to the ways of coping with novelty in animals.

This manuscript has not been published and is not under consideration for publication elsewhere.

## Introduction

Dispersal and population growth are the two fundamental processes that ensure the expansion of populations (Skellam 1951, Phillips and Suarez 2014). Dispersal corresponds to individual movements through space (and into new spaces), and population growth to space filling (including newly colonized space) by individuals. It is generally considered that invasions occur when a species colonizes a habitat that had never been occupied before (Pascal et al. 2003).

Many of the most rapid and famous invasions have involved flying species, such as the House finch, *Carpodacus mexicanus*, the House sparrow *Passer domesticus*, the European starling *Sturnus vulgaris*, the Eurasian Collared dove *Streptopelia decaocto*, and the Gypsy moth, *Lymantria dispar* (Elton 1958, Veit and Lewis 1996). However, flying far away is not enough: dispersers face novel situations and habitats to which their population of origin had never adapted (Sax and Brown 2000). When an invading species spreads, it will face challenges related to novel environments, novel foods and novel objects. Individual variations in neophobia will thus determine which individuals survive, which settle and which do not. Leaving the original colony may also mean some degree of social isolation. Many of the traits associated with invasion are behavioral traits and some of them may be dependent on individual personalities. Thus, genetic or phenotypical variability may also support and explain invasion success (Rejmanek and Richardson 1996, Wilson 1998, Jason et al. 2004, Kolbe et al. 2004).

One classical definition of temperament is that it corresponds to individual behavioral characteristics that are relatively stable over time and across situations (Bates 1989). For Hall et al. (1997), when temperament is refined by experience, it becomes personality. For these authors, the concept of temperament involves some deeply biologically rooted characteristic. This questions the determinism (genetic and/or epigenetic) of these interindividual differences (Digemanse et al. 2002, Hausberger et al. 2004, Van Oers et al. 2004a, Groothuis and Carere 2005). Nowadays, though, both concepts tend to be used interchangeably. While relative stability of traits over time is generally interpreted as reflecting the existence of temperaments or personalities (Jones 1977a, Gosling 2001), its absence is interpreted as the expression of context or state dependent behaviors (Van Oers et al. 2005) or as the expression of behavioral flexibility (i.e. the individual can adapt its behavior to the different situations) (Pfennig et al. 1993, Neff 2003). Although some studies have found evidence of genetic or acquired behavioral phenotypes (Dingemanse and Réale 2005, Pittet et al. 2013) it is generally unclear which individual differences are due to phylogenetic or population history, and which to individual experience (Fox and Millam 2004).

A second important question concerns the constancy of individual differences across situations (Sih et al. 2004). The question here is whether individuals with a particular behavioral response in a situation behave in a particular and systematic way in another situation, hence present “behavioral syndromes” or “coping styles” (Wechsler 1995). There is controversy in the literature with some studies finding stability across situations (Le Scolan et al. 1997, Sih et al. 2003) and others not (Coleman and Wilson 1998, Neff and Sherman 2004, Lee and Tang-Martinez 2009). These different observations may reflect either species differences, differences in the experimental procedures, or both. Habituation and learning are two processes that can also modify behavioral responses in specific contexts.

Studying personalities may be an essential aspect in the understanding of biological invasions, especially neophobia, as stress physiology and behavior are highly relevant in this context (Crane et al. 2020, Greenberg 2003, Martin and Fitzgerald 2005). For example, Atwell et al. (2012) demonstrated that there were rapid adaptive shifts in both stress physiology and positively correlated boldness behavior in a songbird, the dark-eyed junco, *Junco hyemalis* following its colonization of a novel urban environment. They found persistent population differences with both reduced corticosterone responses and bolder exploratory behavior in birds from the colonist population. Furthermore, behavioral flexibility, particularly in relation to novel stimuli and introduction into novel environments, has been suggested as a possible explanation for why some species become successful invaders (Sol et al. 2002, Wright et al. 2010, Webb et al. 2014).

European starlings are well known for their ability to adapt and invade a wide range of habitats, as shown by their expansion in the varied parts of the world where they had been introduced (e.g. Long 1981, Feare 1984, Craig 2020). However, both rural and urban populations can be found. Are these preferred habitats associated with particular individual behavioral characteristics or are they just a result of the availability of nest sites. Also the question arises whether young birds acquire skills to exploit the novel resources and challenges (presence of humans, vehicles…) provided by the urban habitat, or whether heritable population characteristics develop over generations.

In the present study, we hypothesized that more recent urban populations may have adapted to the challenges of urban life and hence that rural and urban young birds would differ in personality traits even when experiencing the same developmental conditions. In order to test this hypothesis, we hand-raised young birds taken from the nest either in rural or urban areas of the same region, where they are sedentary. As young adults, we tested their reactions to novel situations, objects or food, after they had spent one year together under the same conditions. Since migratory populations also face novel situations, we also tested a group of migratory birds wild-caught as adults.

## Methods

### Subjects

Three groups of birds (N=44) were used: two groups of sedentary birds taken from nests in the Brittany region of France, and one group of migratory birds from eastern Europe wild-caught as adults.

The sedentary birds comprised: the “rural group” nestlings hatched within a 20 km radius of Rennes where starlings settled more than 400 years ago (Richard 1826), while the “urban group” hatched within Rennes city, an urban area colonized 30 to 80 years ago (Clergeau 1981).

These 32 nestlings were removed from 32 different nests (one chick per nest to avoid possible sibling effects, either genetic or environmental) when they were 5-14 days old in Spring 2007 (N=18: 12 rural and 6 urban) and 2008 (N= 14, 7 rural and 7 urban). All the young birds were hand-fed using commercial pellets (Végam, Grosset) mixed with water for five weeks until they could eat independently. At the age of two months they were put in an outdoor aviary (3mx4mx2m) as a single group (one for each generation).

Thus, Urban and Rural individuals were always reared together.

<u>The adult migratory group</u> was composed of 12 adult starlings (six males and six females), captured during autumn 2006 with nets at the cliffs of Etretat in Normandy during the migration season. They were at least 2 years old at the time of capture, as estimated by feathers (Feare 1984). They were then housed in a large outdoor aviary with other adult starlings until the beginning of the experiments in autumn 2007.

### Behavioral tests

The birds’ reactions to four different challenging situations were assessed: social separation, novel environment, novel object and novel food. To establish individual responses to familiar food (“baseline”), we also performed a test where the starlings had access to mealworms.

For all the tests we used 120 x 40 x 50 cm cages that were divided into two identical parts (part I and part II) by an opaque plastic barrier. The experimental design and the chronology of the experiments are presented in Figure 1. All tests were videorecorded using a SONY Handycam Digital 8. for later analysis. Thus, the birds were assessed in the following tests:

**Figure 1:**
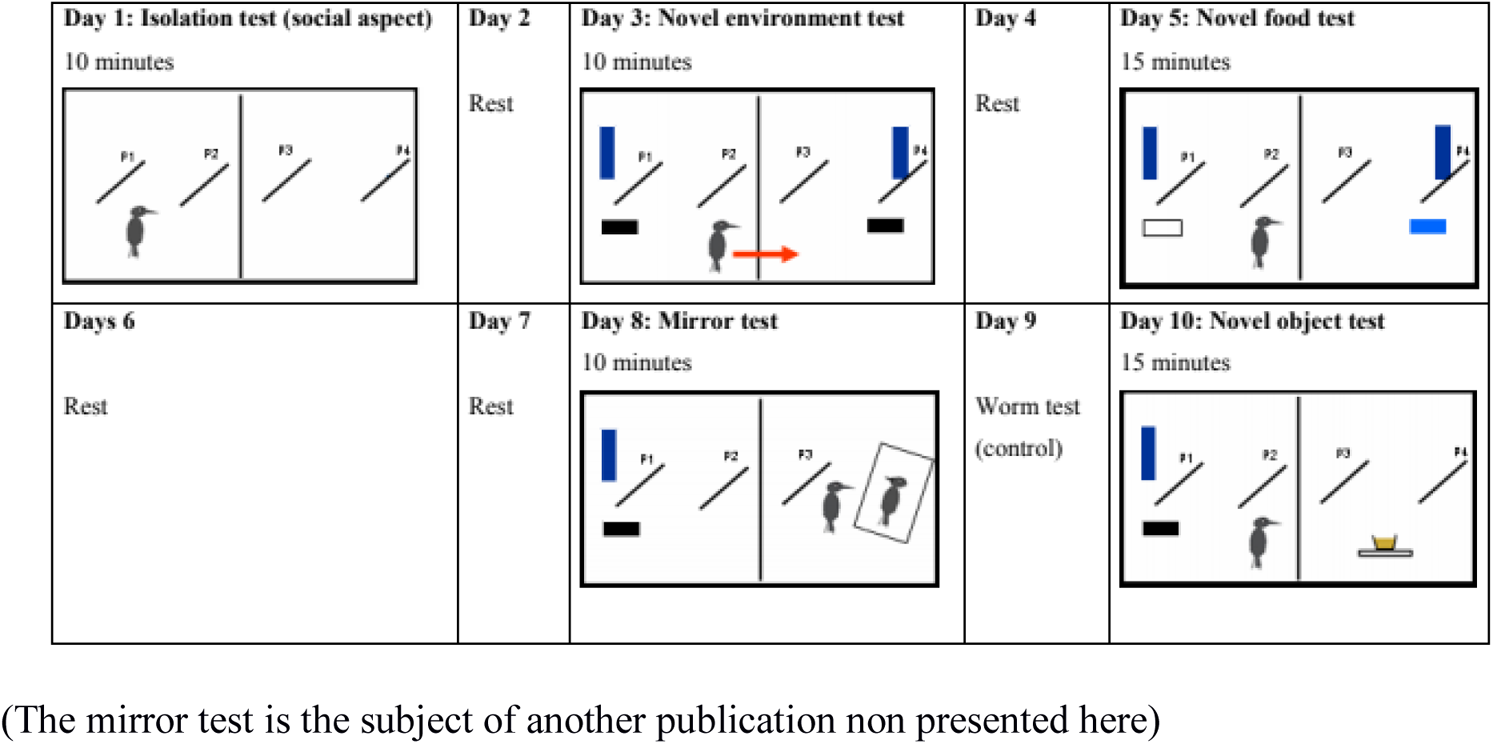
Experimental design and chronology of the experiments conducted on the three groups of starlings.

#### Reaction to social separation

The reactions of the birds when they were first in the individual cages were measured during the 10 first minutes after their arrival. During this part of the experiment there were two perches P1 and P2 in the part I of the cage and no feeding dish or a drinking trough.

#### Novel environment test

After the bird had habituated over 2 days to being in the part 1 of the cage, we removed the plastic barrier so that the birds had access to a larger area. For this test, there were four perches, one feeding dish, and one drinking trough in each part of the cage.

#### Novel and familiar food tests

Food colour is one major factor inducing neophobia in birds (Marples and Roper 1996, Kelly and Marples 2004). Starlings are very sensitive to the colour blue, but this is not a common colour for their usual food (Hart et al. 1998). The novel food consisted of the usual pellets coloured blue using methylene blue (75g commercial food + 50 ml of water + 5ml methylene blue) placed in the usual feeder in part II of the cage. The test lasted 15 minutes.

The individuals were deprived of standard food 30 minutes before the experiment, in order to increase their food motivation.

One test was also performed using familiar food in order to establish individual responses to food: three mealworms were placed in the feeding dish in part II of the cage and the latencies to approach the feeding dish and eat mealworms were measured. This test was conducted four days after the blue food test, and one day before the novel object test.

#### Novel object test

A novel object (a Petri dish wrapped in clear tape and attached to a white wood substrate) was placed in part II while the bird was in part I (closed). The barrier was then removed so that the bird could see and approach the object. The test lasted 15 minutes.

### Data analysis

Behavioral data were analysed using continuous focal sampling (Altman 1974). The list of behaviors recorded is in Table 1.

**Table 1:**
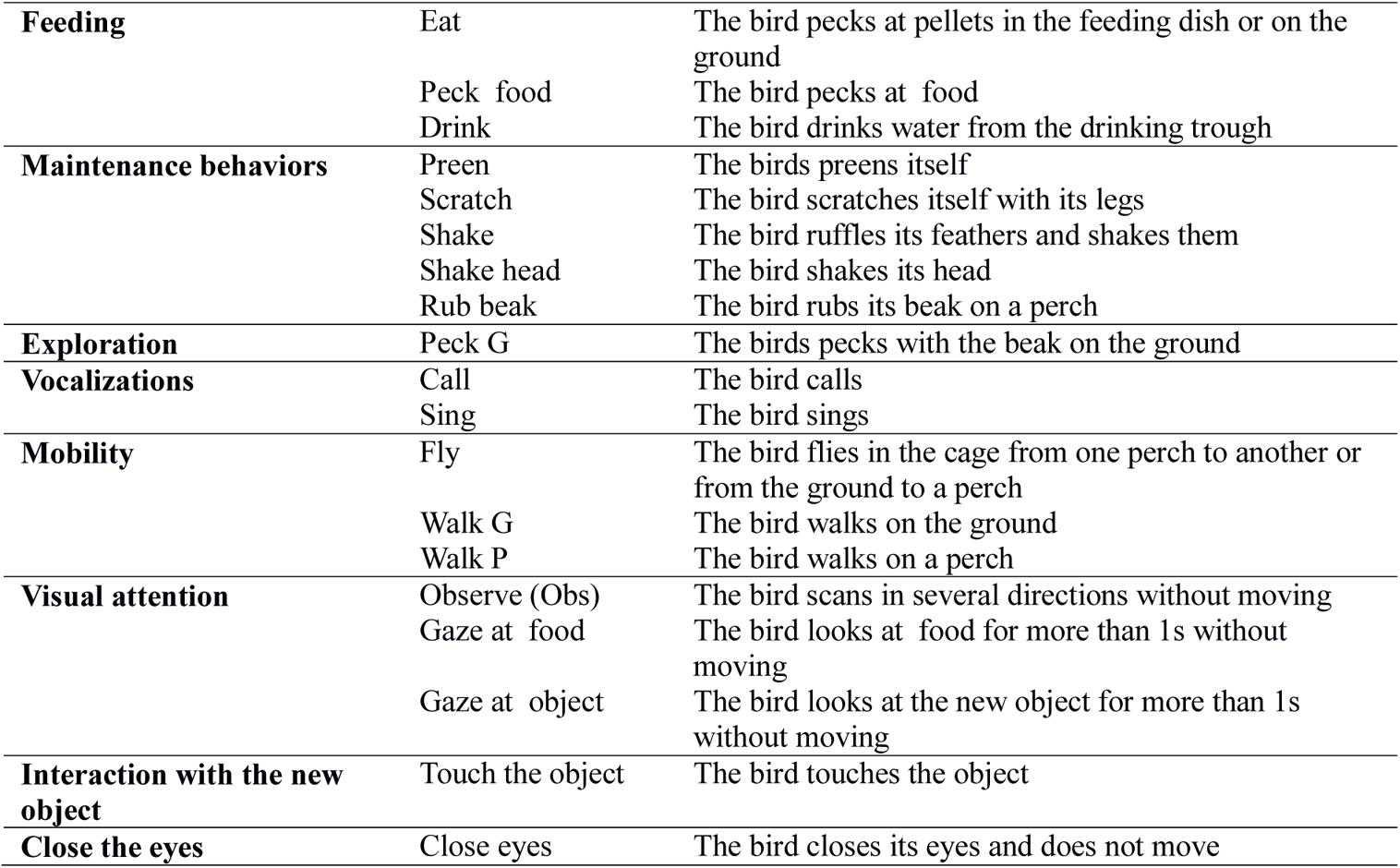
Behaviors recorded during the isolation and neophobia tests.

Temporal parameters were also measured such as the latency to enter part II in the novel environment tests, to peck at the novel food or to approach (less than 20 cm) or touch the novel object in the food and novel object tests respectively.

For the novel food test, the number of pecks at the food and the weight consumed were also measured (feeding dish weighed before and after the experiment).

For the novel object test, the number of times the bird touched the object with its body (with its legs or its beak) was also taken into account.

#### Statistical analysis

We used Cox models implemented in R 2.8.1 software to compare the probability of approaching the new situations between the different groups, and to test for potential effects of sex and year of capture on these probabilities (Cox and Oakes, 1984).

As there were no effects of sex and year of capture on behavior, we grouped the data of males and females and of the different years of capture in each category (rural and urban young).

Given the sample size, normality was not ensured and non-parametric tests (Kruskal-Wallis and Mann-Whitney) were used to compare the groups of birds (Siegel 1956).

Pearson R correlation coefficients were calculated to test for potential correlations between the different parameters measured.

#### Behavioral profiles

For the first three tests, hierarchical ascendant classifications were performed using R 2.8.1 software and the Ward method of clustering in order to group individuals with behavioral similarities and detect different behavioral profiles (Ward 1963).

For each test, the analysis was performed on the basis of specific measures:

- Isolation test, eight behaviors were recorded: Walk on the perch, Walk on the ground, Fly, Pecks on the ground, Vigilance, Call, Maintenance.
- Novel environment test, four were used : Walking on the ground and Flying, as well as the time required to enter the novel environment and Time spent in the novel environment. The maximum time was 600 seconds (the duration of the test).
- Novel food test, five behavioral parameters were used: frequencies of flying and gazing at the food, time to taste blue food, number of pecks at food, and quantity of consumed food. When the individuals did not taste the blue food, the value used for the latency was 900 seconds (the duration of the test).

Once we had obtained the different clusters for each experiment, we conducted Kruskal-Wallis and Mann-Whitney non-parametrical tests between the clusters in order to detect which behavioral items distinguished them.

Finally, we conducted Chi square tests on the number of birds from each group (rural, urban or migratory) in order to see if there were significant differences between the groups in the proportions of individuals presenting each profile.

For the novel object test, we compared the proportion of individuals that approached the object and the proportion of individuals that touched the object in each group.

#### Comparison of birds’ reactions between situations

Pearson R coefficient tests were performed in order to test if there were correlations between latencies across the tests:

- to taste the blue food versus time to enter the novel environment
- to taste the blue food versus time to eat the first worm
- to eat the first worm versus time to enter the novel environment
- to approach the novel object versus time to enter the novel environment
- to approach the novel object versus time to taste the blue food

We conducted Kendall correlation tests in order to measure the degree of correspondence between behavioral ranks across the tests. For example, the individual ranks of the frequency of flying in the novel environment test were compared to the ranks of flying in the novel food test.

### Animal welfare note

This study was conducted at the Ethos Laboratory at Rennes University approved by the French Comittee for Animal Welfare and the French Ministry of Research, following the recommendations for taking care of and experimenting on Starlings. The European starling is an invasive non protected species. Furthermore the present study was based on behavioral observations and was strictly non invasive in physical terms.

## Results

### Behavioral responses

There was no effect of the year of capture, age or sex on the behavioral responses of the birds in the different tests (Cox analyses: 1.13<OR<1.63 0.192<p<0.8 OR=Odd Ratio) However clear differences appeared between the two groups of hand-raised birds in the novel environment test OR=2.5 p=0.035). Twice more urban birds entered the novel environment during the test than rural birds(Cox analysis, confidence interval = [1.07; 5.85] (Figure 2).

**Figure 2:**
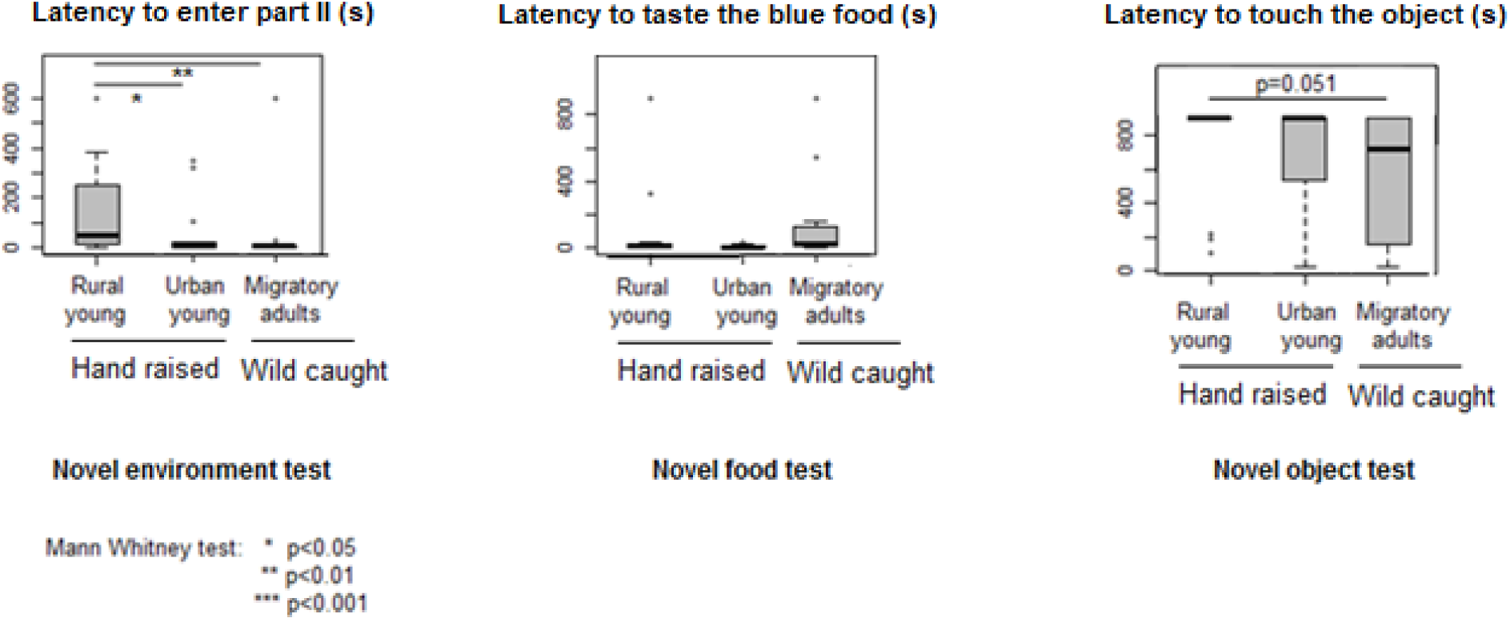
Comparison of reactions to neophobia tests between the three groups.

There was a significant difference in the latency to enter the novel environment between the three groups (Kruskal Wallis test: H=10.45, p=0.0055) but none for the other latencies (p>0.05).

The urban and migratory birds entered the novel environment more quickly than the rural birds (Xr= 176sec, Xu= 67sec, Nr=19, Nu=13, z=-2.19, p=0.03, Xm= 57sec Nm=12 z=-3.11, p=0.0019<0.017) while there was no significant difference between the urban and migratory birds (z=-0.44, p=0.66) (Figure 2).

The rural birds tended to touch the object later than the migratory birds (z=153, p=0.051) (Nr=19, Nm=12, Xr=532sec, Xm=418sec).

There was high interindividual variability in the latencies to enter part II (1 to 600 seconds) and in the time spent there (0 to 594 sec.), but both times were negatively correlated (Pearson test: R=0.7, df=42, p<0.05).

Similarly, there was a large diversity of reactions in the novel food test, latencies to peck at the blue food range from between 1 to > 900s as some individuals never tasted the food during the 15 minutes of the test.

The number of pecks to the food comprised between 0 and 187.

When we conducted Pearson tests, there was a negative correlation between the latency to taste the blue food and the number of pecks to it (R=0.33 df=42 p<0.05) indicating that the earlier the birds tasted it the more they subsequently ate, as also shown by the correlation between the number of pecks at food and the weight consumed (R=0.61 df=42 p<0.01, 0 to 3.03g).

Eighteen birds never approached the novel object while others did so within 3 seconds. There was no correlation between the latency to approach and the latency to touch the object (R=0.29 df=25 p>0.05) nor between the number of contacts with the object and the latency to touch it (R=0.288 df=24 p>0.05).

### Behavioral profiles

#### Social isolation test

The hierarchical ascendant classification lead to the identification of two different behavioral profiles: a calm profile and an active profile (Figure 3). Individuals from the active profile were characterized by high levels of mobility and showed the following behaviors significantly more frequently: flying, walking on the perches, walking on the ground, pecking on the ground, observation and distress calls (Figure 4). The individuals in the calm group tended to show more maintenance behaviors.

**Figure 3:**
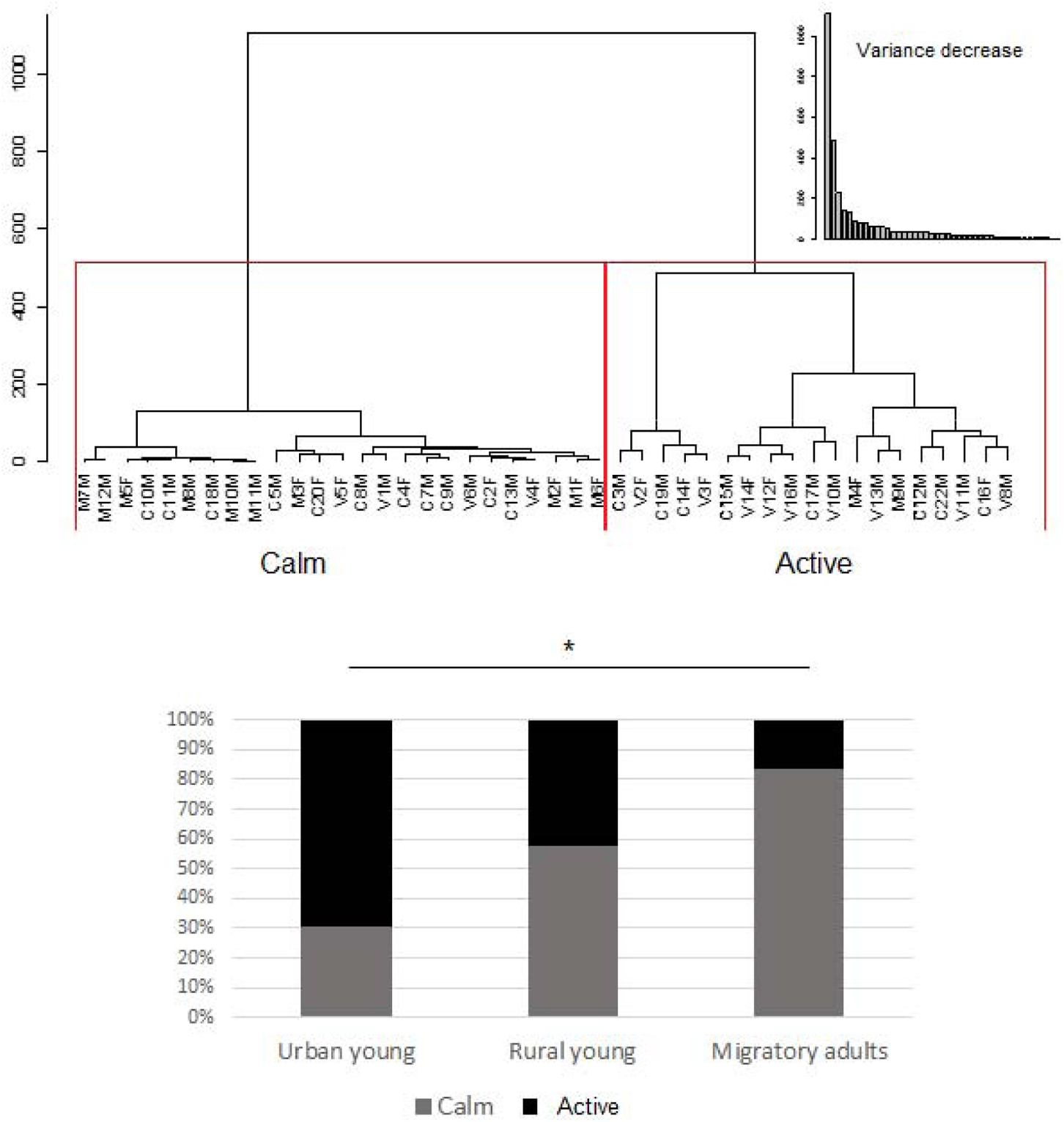
Clusters from the hierarchical ascendant analysis on isolation test (C=Rural, V=Urban, M=Migratory)

**Figure 4:**
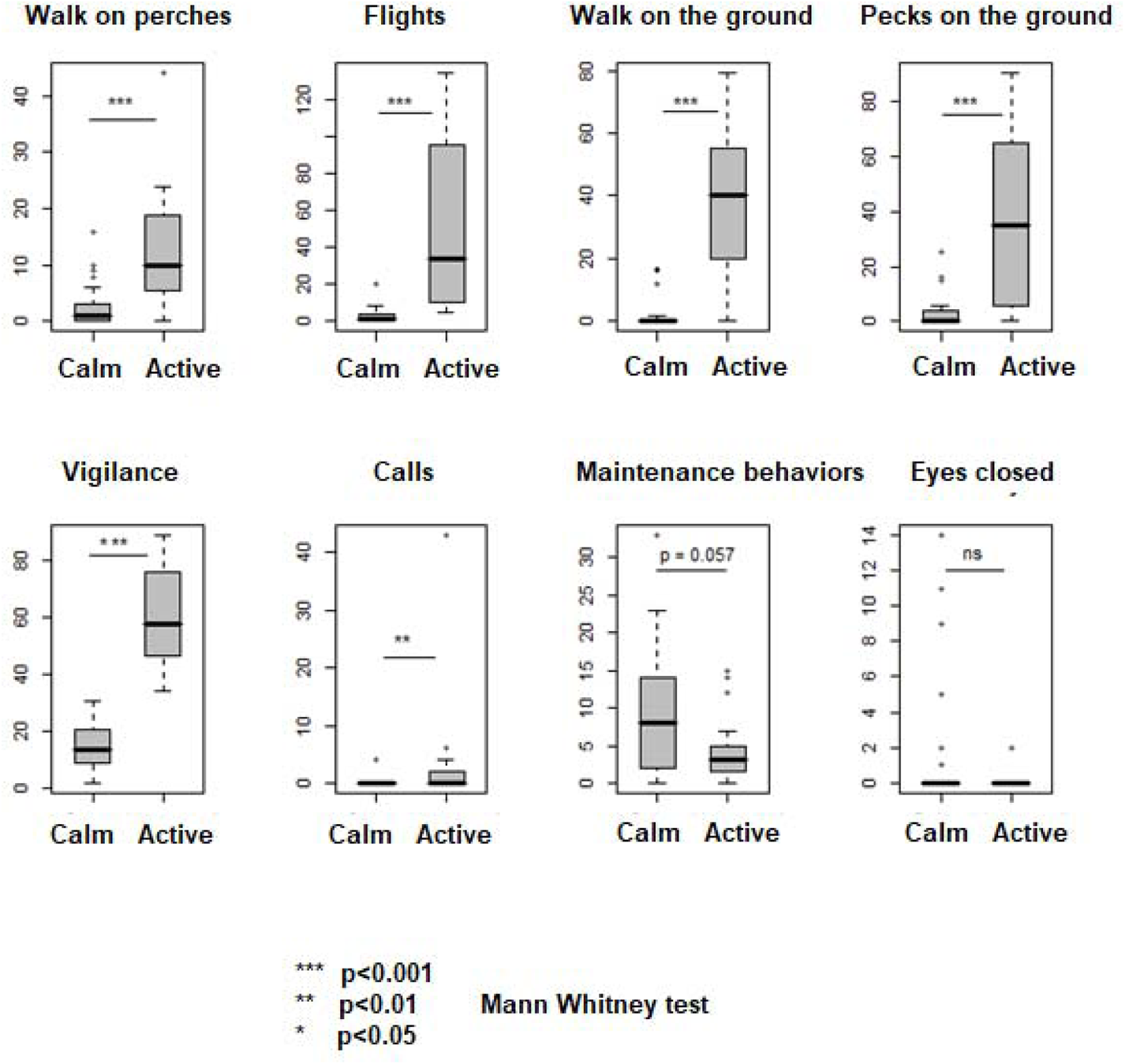
Behaviors expressed by each cluster in the isolation test.

The frequencies of individuals in each group indicated that there were significantly more active individuals in the urban young group than in the migratory adult group (χ p<0.05)

#### Novel environment test

The hierarchical ascendant classification allowed us to distinguish three different behavioral profiles (Figure 5): a shy profile, an intermediate profile and a bold one.

**Figure 5:**
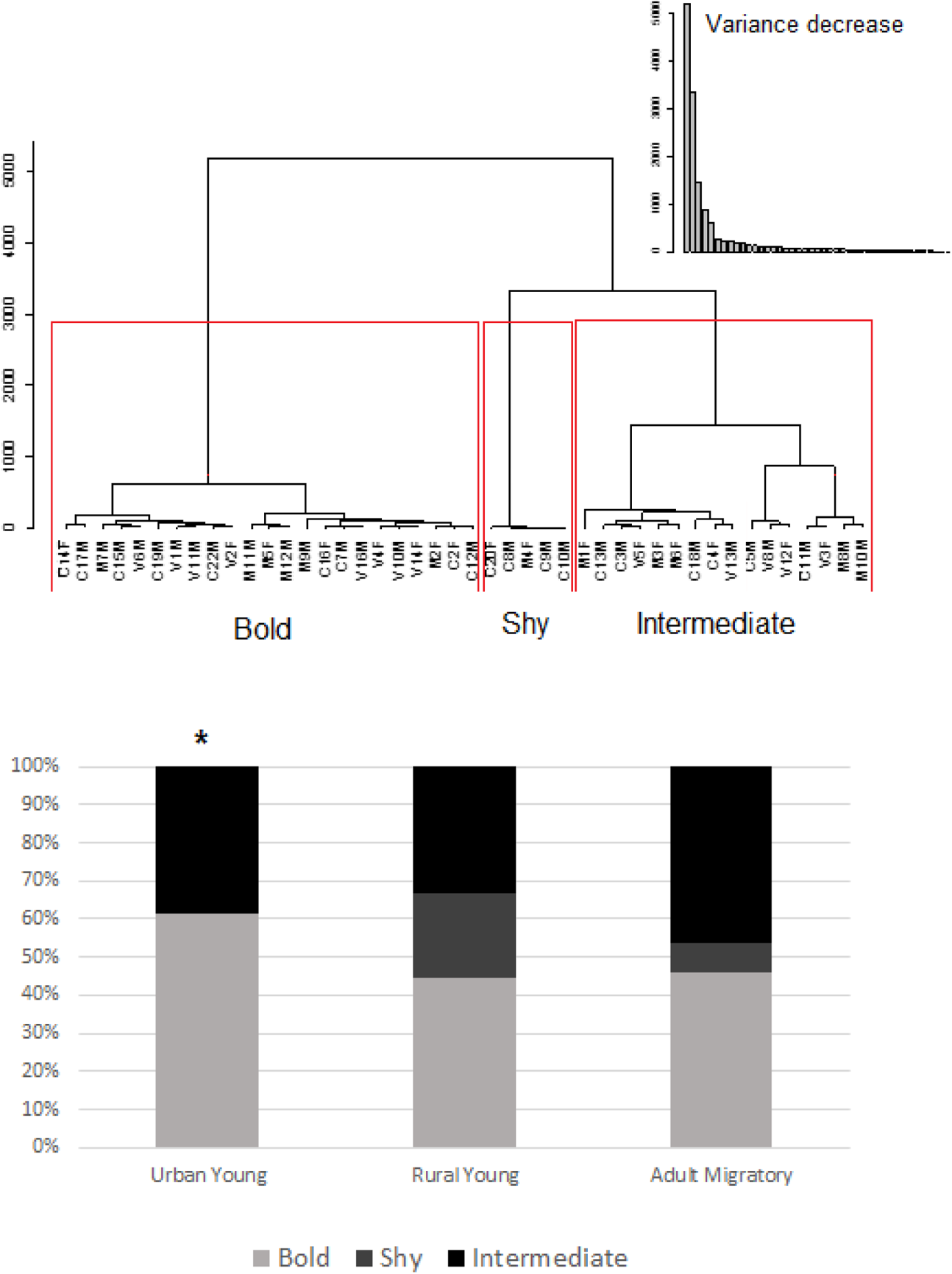
Clusters from the hierarchical ascendant analysis of the novel environment test.

The bold and the intermediate profiles differed significantly from the shy profile in many ways: higher levels of mobility (walk and flights), and birds entered the novel environment while the shy individuals did not(Figure 6). Bold and intermediate individuals differed in the time they spent in the second part of the cage: the bold ones stayed more than half of the time of the experiment in the second part of the cage, significantly longer than the intermediate ones that stayed less than 300 seconds in the novel environment.

**Figure 6:**
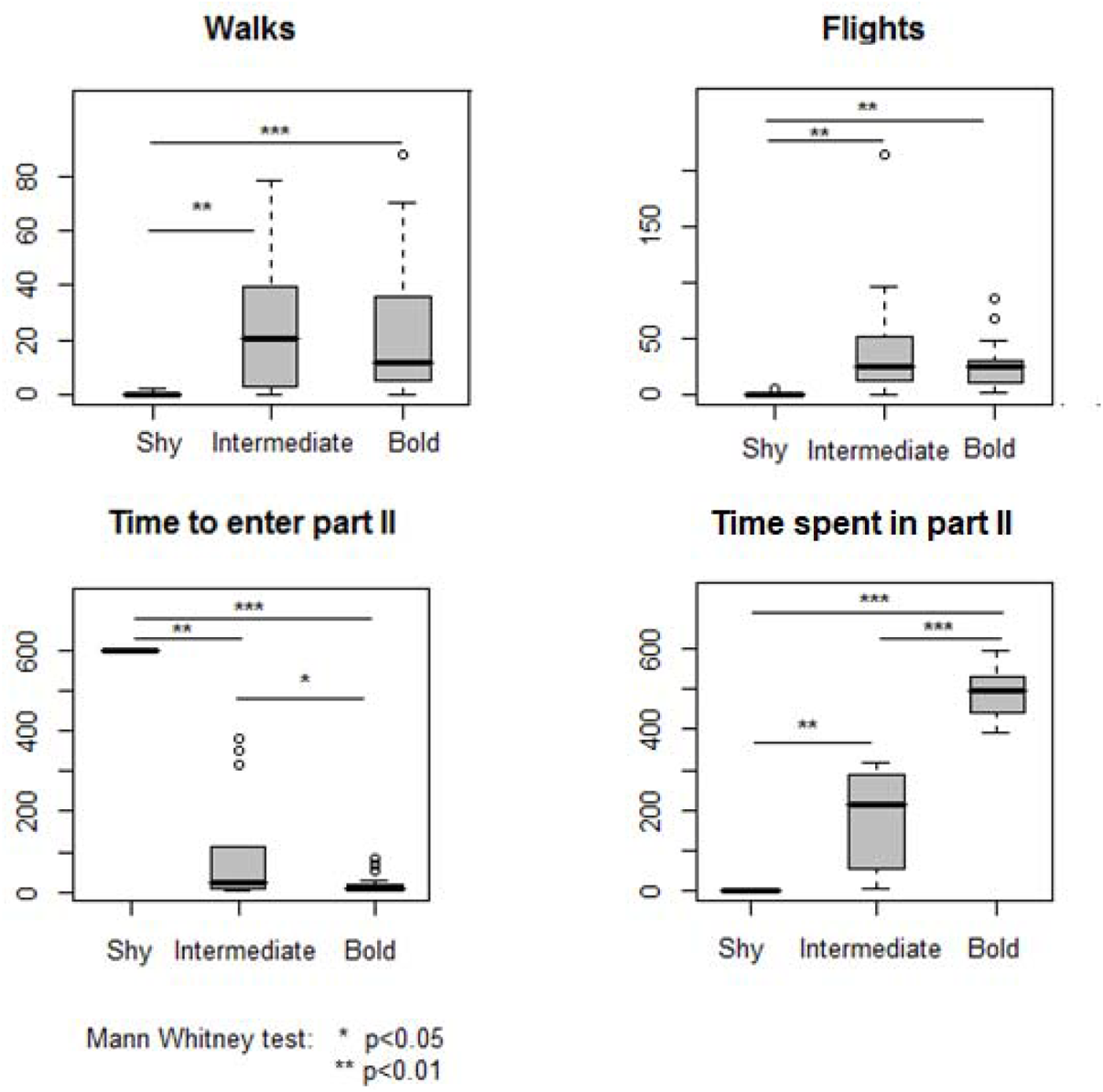
Behaviors expressed by each cluster in the novel environment test.

There were significantly more bold individuals than shy ones in the group of urban birds (Χ² test : p=0.023). The proportions of bold, intermediate and shy individuals were equivalent in rural and migratory groups (Χ² test : p<0.05).

#### Novel food test

Three different types of profiles appeared again: a bold, a shy and an intermediate profiles (Figure 7). The latency to taste the food was longer in the shy individuals (Figure 8). The bold birds pecked significantly more at the food and ate more of it than the intermediate birds (Figure 8). These two profiles ate more than the shy individuals who very rarely pecked at the food. The proportions of the different profiles did not differ between the three groups and nor within each group (χ² test: p > 0.05).

**Figure 7:**
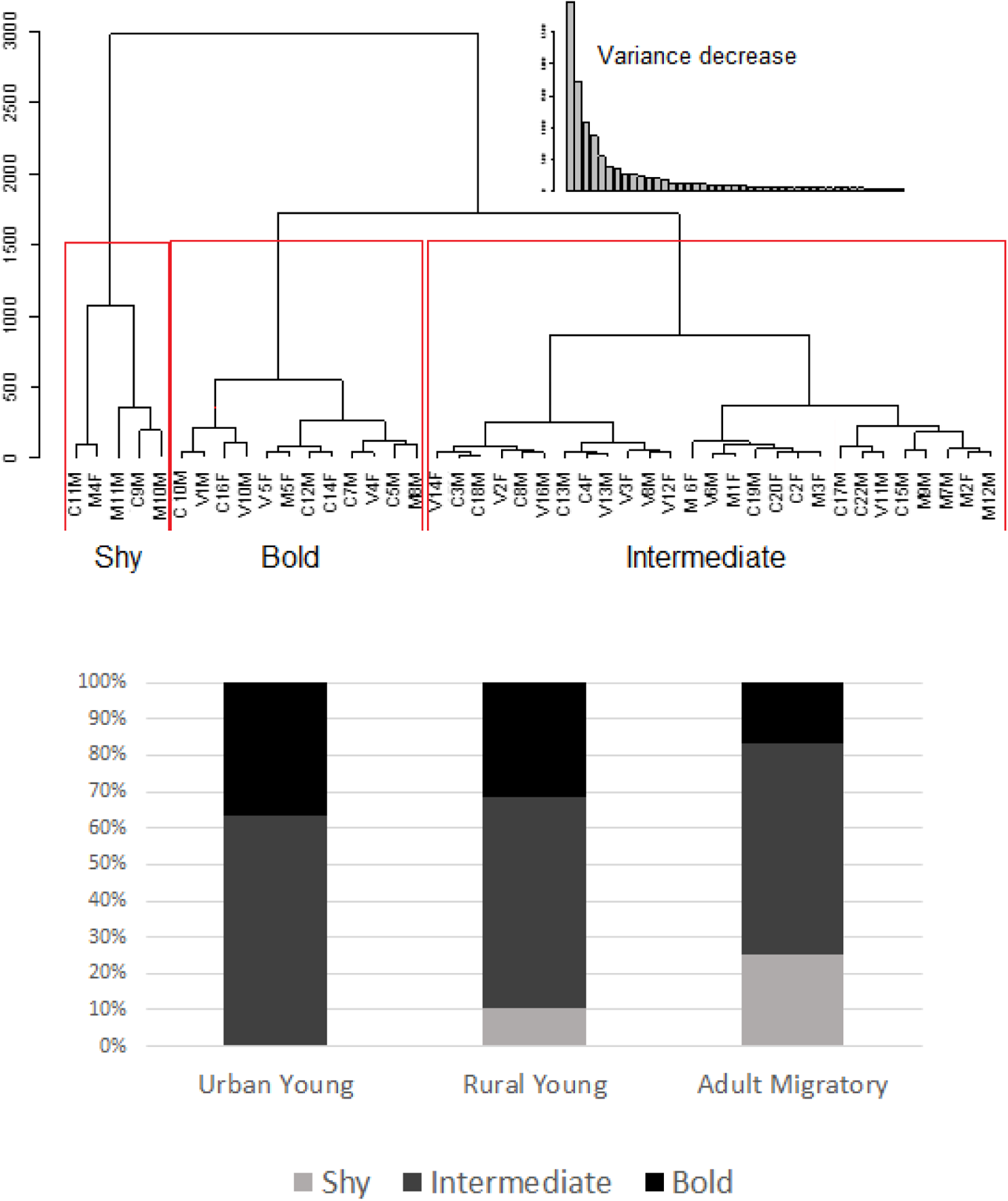
Clusters from the hierarchical ascendant analysis on novel food test.

**Figure 8:**
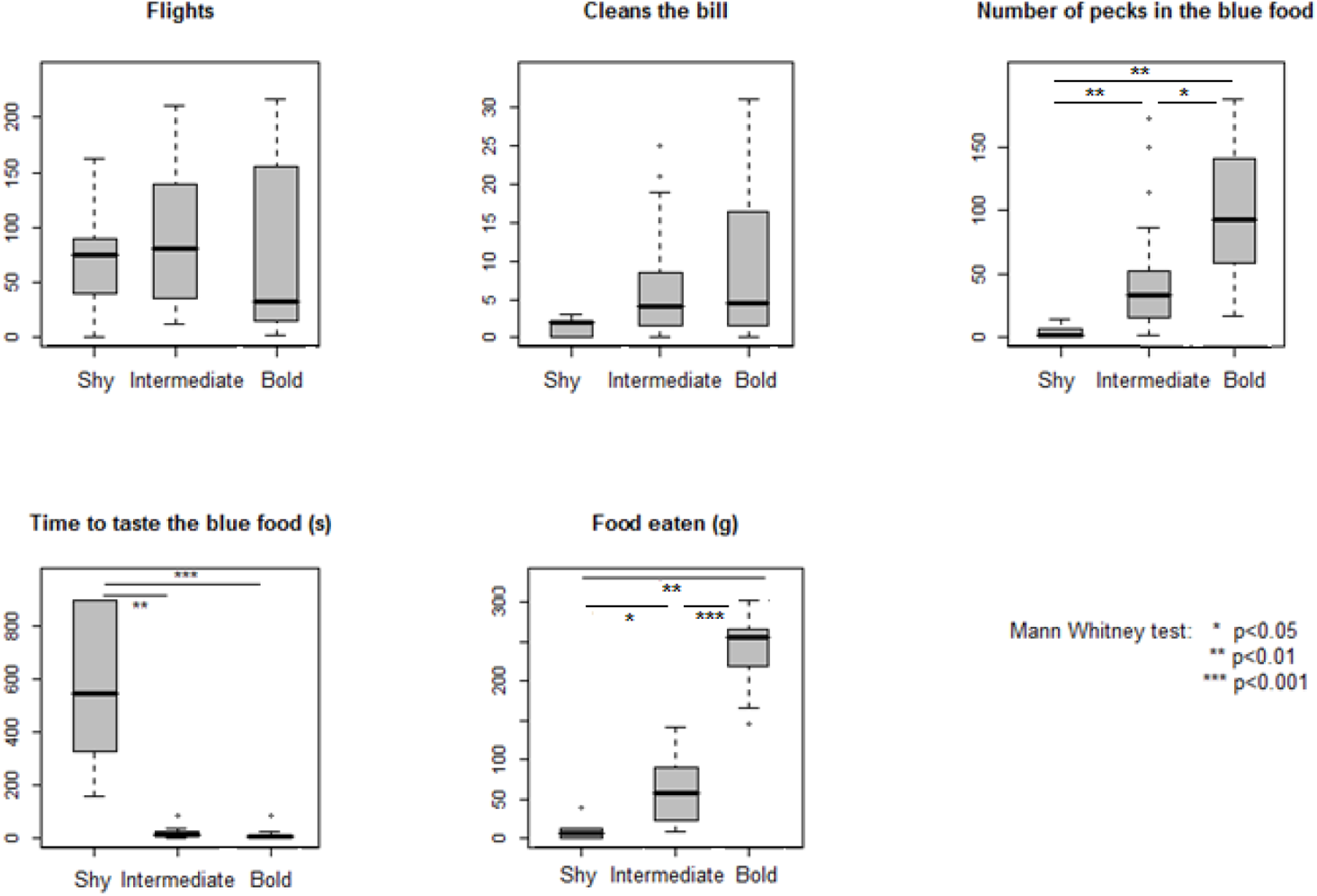
Behaviors expressed by each cluster in the novel food test.

#### Novel object test

As mentioned earlier, a large proportion of birds never approached or touched the object. The proportion of these clearly neophobic animals did not differ between groups p>0.05 for χ² tests.

However, in the rural young group there were significantly more individuals that did not touch the object than individuals that did (Figure 9).

**Figure 9:**
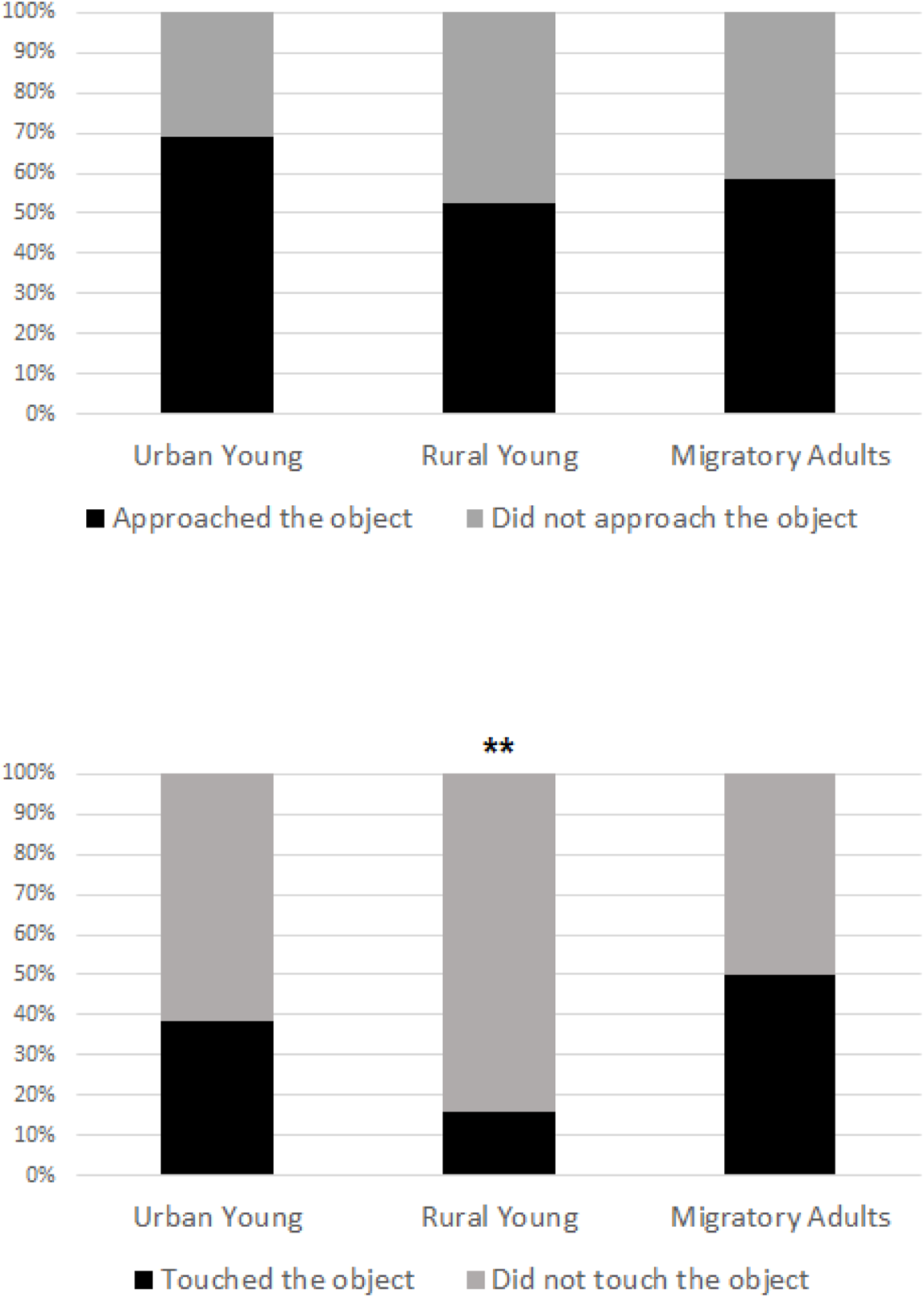
Proportions of individuals that approached and that touched the object.

### Correlations between tests

There were no correlations between the latencies (to enter the second part of the cage, to taste the blue food, to approach the object) observed in the three neophobia tests (0.0057< R < 0.29 df=42: p>0.05).

There was a significant positive correlation between the latency to test the blue food and the latency to eat the first worm (R=0.56 df=42 p<0.01).

### Correlations in behavioral ranks between tests

Some behaviors were correlated between tests (Table 2):

- walks on perches, pecks on the ground and vigilance frequencies between the isolation and the novel environment tests
- frequencies of walking on the perch, flight, walking on the ground, observation, calling; respectively between the isolation and the novel food tests (i.e. 5 out of 7 behaviors measured), frequencies of walking, flying, observing and calling; respectively between the isolation and novel object tests
- the number of walks on the perches and the rank of visual attention; respectively between the novel environment and the novel food tests,
- the ranks for the number of walks, the number of times the birds fed and the ranks of vigilance frequency between the novel object and novel environment tests,
- the ranks of the number of walks on the perch, the ranks for visual attention (vigilance behaviors and gazing at the novel item, food or object) between the novel food and novel object tests.
- the ranks to eat food between the novel environment and the novel object tests, indicating that the birds ate similar quantities when the food had the same familiar aspect. However, the ranks to eat blue food were not correlated to the ranks to eat the normal non-coloured food indicating that the change of colour alters the usual levels of consumption of birds.

**Table 2:**
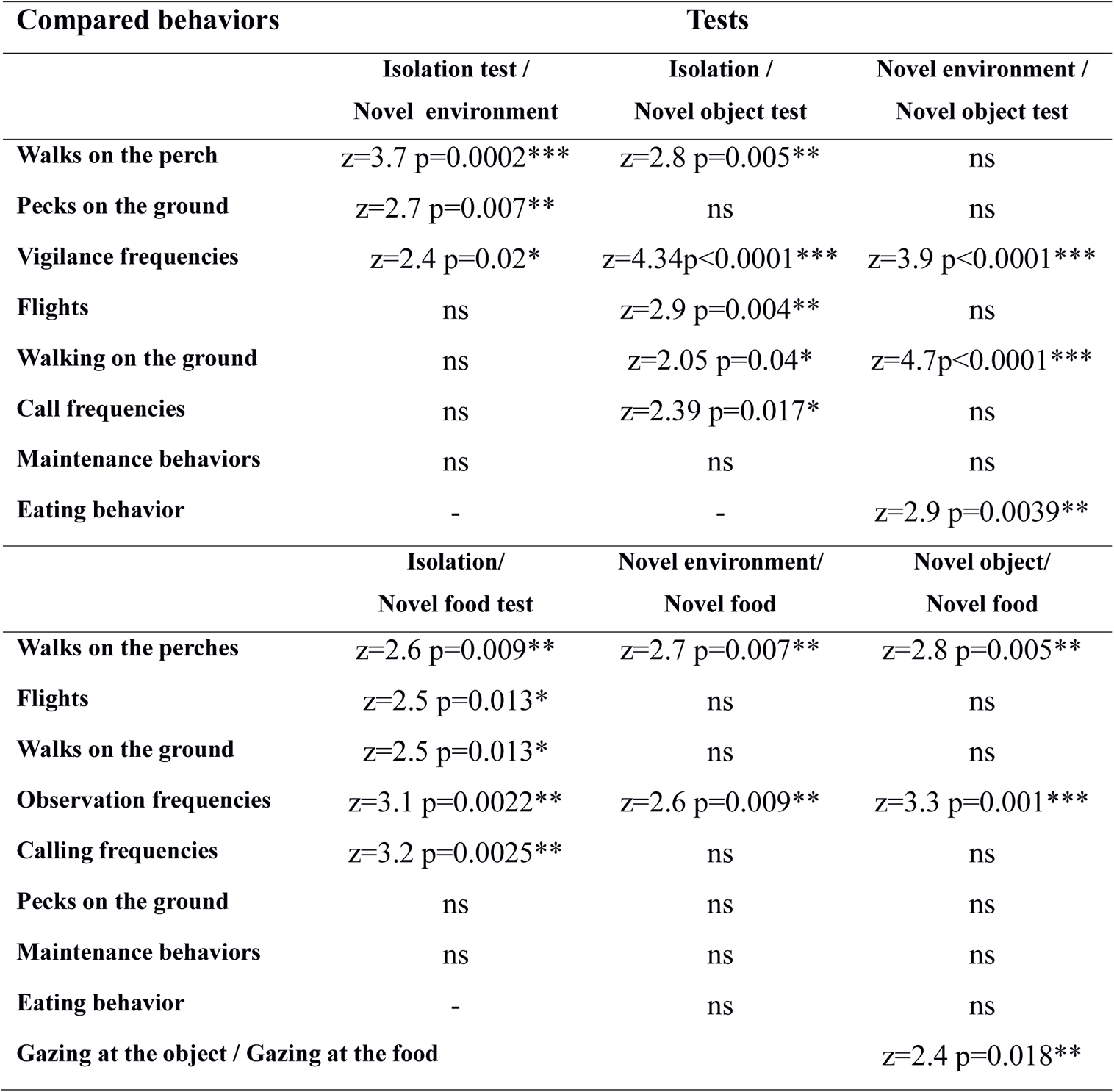
Consistencies in birds’ behaviors between tests: p values for the correlation of ranks in the Kendal test.

### Modifications of behavioral responses between tests

The comparison of the frequency of behaviors between the two tests with the same duration (novel environment and novel object test) revealed that the birds performed more flights in the novel object test than in the novel environment test, suggesting a higher level of fear when they were faced with a new object (Wilcoxon test, N=44, z=-4.438; p<0.0001).

For the other behaviors, the differences were not significant (p>0.5 in all cases).

## Discussion

This study, based on behavioral tests, reveals that different invasion histories of previous bird generations were reflected in personality differences in current generations: even when hand-raised and maintained together under similar conditions, starlings from a rural non-invasive population proved more reluctant to touch a novel object and to enter a novel environment than those from an urban invasive population. The comparison with migratory birds caught as adults revealed that they showed more similarities with the urban birds. Careful examination of individual behavioral profiles produced clear groups that could differ according to the trait tested. Thus, two profiles (calm/active) emerged in the social separation test, and three profiles (shy, intermediate and bold) in the novel environment and the novel food tests.

Some individual stability, as shown by correlations between tests, suggests that these were indeed individual stable behavioral differences.

The different populations differed in their representation within each personality cluster. Overall, the rural non-invasive birds appeared calmer in the social separation test but shyer in all neophobia tests than the urban invasive birds, even though they had shared the same developmental history (ontogeny). This probably indicates that behavioral traits that may have characterized their ancestors have been inherited and retained.

The migratory birds, caught as adults, appeared to have an intermediate profile, or even tended to be shyer, which may be partly explained by their quite different life history.

### Social profiles in an invasive bird

In the isolation test, we observed two types of reaction (active and calm). The active individuals are probably more emotive and more stressed by social deprivation. Calm and active individuals were observed in all three groups. However, the urban group contained a higher proportion of active individuals in comparison to the migratory group.

These results match our observations in the field where we found that individuals from colonisation fronts are more readily attracted to decoys and to starling song playbacks (in particular in recently colonized urban areas) (Rodriguez et al. 2010a, Rodriguez et al. 2020). These populations seem to comprise more individuals who actively seek social contact. Nevertheless, leaving a colony to settle in an unoccupied habitat may favour calm individuals that are tolerant of social isolation at colonization fronts. During field observations in southern Italy, one of the more recent propagation fronts of the species in Europe, we found one pair of starlings nesting alone in the rural area of Otranto (Rodriguez 2010b). This pair was five kilometres from the nearest other pairs during two consecutive years. Hausberger (1986, 1988) also observed isolated pairs breeding on the colonisation front of the Australian invasive population. Breeding in social isolation is thus not impossible for this usually gregarious species.

To our knowledge this is the first time that two types of reaction to social isolation have been described in an invasive species. We suggest that the existence of these two different strategies enables behavioral flexibility in situations with different population densities. Such flexibility has been demonstrated in the vocal communication of European starlings which differs according to population density (Henry et al. 2015). Similarly in yellow-bellied marmots (*Marmota flaviventris*) females that had affiliative interactions with more individuals, and those that were more socially embedded in their groups, were less likely to disperse (Blumstein et al. 2009). Fogarty et al. (2011) found, by modeling invasive processes, that expansion occurs more rapidly when a species has a mix of life-history or personality types that differ in density-dependent performance and dispersal tendencies. They also found that polymorphism in sociability increases the rate of advance of the invasion front, since asocial individuals colonize empty patches and facilitate the local growth of social types that, in turn, induce faster dispersal of “asocial” individuals at the invasion edge. Our results are in agreement with this model as we found different kinds of reactions towards social isolation indicating a mix of personalities in the first test.

### Neophobia and exploration in invasive processes

When individuals leave their habitat of origin during migration or when they settle in new habitats, they are likely to encounter new food items or novel objects. All groups (rural young, urban young and adult migratory) included some individuals who entered the novel environment rapidly, and stayed there for a long time apparently at ease, indicating occupying novel environments without expressions of stress is not restricted to one particular group.

The analysis of starling behavioral responses to neophobia revealed a high diversity of reactions towards novel situations in the species. These interindividual differences were observed both in latencies to approach novelty and in behavioral profiles indicating that the species contains a wide range of possible responses to cope with novel environments. We observed a continuum of responses from shy to bold reactions with intermediate levels of mobility and of latencies to approach novelty.

### Neophobia towards novel environments and consequences for dispersal

The proportion of bold individuals was higher than the proportion of shy individuals in the urban young group. Both urban young and migratory adults entered the novel environment sooner than the rural young did.

We hypothesized that individuals at colonization fronts and migratory birds are less reluctant to explore novel habitats and we found some support for this hypothesis. More studies should be conducted in the field to verify if bold individuals disperse more.

At the intraspecific scale, a relationship between neophobia and dispersal has been observed in Great tits, *Parus major (*Digenmanse et al. 2003), and in a terrestrial tortoise, *Testudo hermanni (*Rodriguez et al. 2018). Individuals that appeared bold or very mobile in neophobia tests travelled greater distances than shy ones when released into the wild (Digenmanse et al. 2003, Rodriguez et al. 2018). Moreover, Dingemanse et al. (2003) observed that great tits assessed as bold had offspring that dispersed further. They also found that immigrant individuals arriving in a new habitat explored novel environments more rapidly than locally born individuals in laboratory tests.

At the interspecific scale, Rehage and Sih (2004) had observed dispersal differences in latencies to leave the original pool and enter new pools in four fish species: *Gambusia holbrooki, Gambusia affinis, Gambusia geiseri and Gambusia hispaniolae*. The two invasive species showed lower latencies compared to two non-invasive species.

Low neophobia towards novel environments can thus probably enhance dispersal and invasion of new habitats but the limits of our experimental design cannot confirm this aspect.

### Neophobia towards novel food

Individual differences in the consumption of novel food were also observed: some individuals did not taste the blue food, some individuals tasted it rapidly but did not eat it again or only a few times while others tasted it rapidly and consumed a lot of it. There were no differences between groups but each group contained bold individuals who took unfamiliar food early and ate large amounts of it.

Rennes city has been reported as a sub-optimal habitat for starlings, where finding sufficient food for nestlings is particularly difficult and higher rates of nestling mortality occur in this population than in the surrounding rural areas (Mennechez and Clergeau 2006). Therefore, the inclusion of novel food items in their diet can be more important for urban than for rural populations in Brittany. Martin and Fitzgerald (2005) have observed that invading house sparrows on the propagation front approached and consumed novel food faster than individuals from areas where sparrows have been settled for a long time.

Here we observed neophilic birds in each group. The phenomenon of neophilia has been reported in many social species of birds, primates and rodents (Galef 1993, Visalbergi and Fragazi 1994, Cadieu et al. 1995, Wauters et al. 2002) where only a few bold individuals take the first step, and then others copy the choices made by bolder ones. Individuals who readily sample novel items and consume large quantities of novel food can expand their diet, whereas individuals who taste only small quantities may detect possible harmful items (Galef 1993). Finally, individuals that do not taste novel food can avoid the consumption of toxic items (Galef and Laland 2005).

In invasion processes, neophilic individuals are probably responsible for feeding innovations. At the colonisation front in southern Italy we observed starlings feeding their chicks with olives and dates, two food items not consumed by older established populations in northern Italy and Europe. Feeding young with novel food may serve to introduce food innovations into the diet of invasive populations.

### Neophobia towards novel objects

Starlings seemed more reluctant to approach novel objects than the new environment or the new food, probably because it is a more artificial (less usual) situation in natural conditions. They appeared to be more fearful as they flew more often in this test.

Nevertheless, half of the individuals of each group approached the object (even in the absence of a food motivation).

Significantly more individuals did not touch the object in the rural young group. In the urban group and the migratory group, half of the individuals were shy while the others behaved in a bolder way and touched the object.

The higher proportion of individuals who touched the object in urban and migratory group is probably due to a longer history of populations encountering novel objects on migration and in urban contexts. Such individuals would probably touch and manipulate objects more readily in the wild. However, observations of free-living birds are needed, as the restrained situation in captivity obliges birds to be near objects that they could ignore in the wild (Greenberg 2003).

Echeverria et al (2006) have observed that when exposed to novel objects close to feeders, birds from various urban and non-urban species expressed neophobic behavior were reluctant to approach. Only the Chalk-browed Mockingbird *Mimus saturninus* expressed a neophilic response towards the novel objects. The authors made their observations in suburban areas with few novel objects. They suggested that urban birds could have become less neophobic by the experience of frequent encounters with novel objects. Lower levels of neophobia were reported in migratory garden warbler, *Sylvia borin*, when compared to resident Sardinian warblers, *Sylvia melanocephala momus* (a non-migratory species) (Mettke-Hofmann et al. 2005). We compared different populations from a species containing both migratory and sedentary individuals, and found a mix of both neophilic and neophobic individuals in the different groups with predominantly neophobic individuals in the sedentary rural group. In a study conducted by Candler and Bernal (2015), differences of boldness were observed in cane toads where individuals from native populations did not approach a novel object while more than half of the individuals from introduced populations did. It has also been reported that early experience with novel objects in laboratory environments can result in low neophobia levels in young hand reared parrots *Amazona amazonica* compared with individuals raised by their parents in simple nest box environments with a lower diversity of objects (Fox and Millam 2004). However, when faced with predator-like images, European starlings hand-reared in the laboratory appeared more reactive than birds wild-caught as adults (Belin et al. 2018). Moreover, there were both differences between our migratory birds and both hand-raised groups and similarities between the migratory wild-caught birds in some respects and the urban hand-reared birds in other respects. Early experience with an “enriched” environment therefore cannot be the sole explanation.

### Possible determinism of personality differences

Differences in personality between starlings may be due to genetic or environmental causes. For example, for the migratory group wild-caught as adults, migration experience may play a role and for the hand-reared birds, developmental experience may have an influence on behavior.

We can imagine scenarios in which successive selection of bold individuals can operate (that means that genetically and physiologically conditioned individuals are selected in colonization fronts by natural processes) in the same way as selection for calm individuals operates in domestication processes (Belyaev 1978). Early parental effects can also play a role. In the quail *Coturnix coturnix,* chicks raised by experienced females are less fearful than those raised by naïve breeders. The less fearful chicks are quicker to explore an area containing a novel object (Pittet et al. 2013).

Female quails submitted to a stress condition lay eggs that contain higher levels of yolk testosterone and their chicks are more fearful in the novel environment test than chicks from females who were not stressed (Guibert et al. 2011).

### Behavioral syndromes in Starlings?

The ranking of flight behavior, visual attention, calling and walking frequencies were positively correlated between the tests, indicating that individual differences are maintained in different novel contexts. Lee and Tang-Martinez (2009) found that in prairie voles the latencies to approach novelty were correlated between experiments involving the same kind of novelty but not between contexts involving really different situations. In horses, whereas there is a correlation between assessments of emotional reactions in similar situations (e.g. social isolation), no correlation was found between different tests (novel object/ novel obstacle), which reflected different interplays between genetic and environmental factors (Le Scolan et al 1997, Hausberger et al 2004).The individual stability in the reaction types observed here probably reflects behavioral syndromes. It would be interesting to conduct physiological analyses to compare bold and shy individuals in different situations.

### Similarity between sexes

We did not observe differences between sexes in latencies to approach the novel situations in any of the three neophobia tests. These observations are in agreement with the results of studies conducted in other species like cats and great tits (Durr and Smith 1997, Van Oers et al. 2004b), but differed from other studies. Jones (1977b, 1982, 1986) observed that female chicks fed significantly sooner, longer and more than males when presented with novel blue food and that females showed less behavioral inhibition when placed in a novel environment or in an open field. In the same way, female rodents seem to explore a novel environment sooner than males (Gray 1971). In other mammals like primates females may be more fearful than males, whereas there are no sex difference in horses (Buirski et al. 1978, Crepeau and Newman 1991, Hausberger et al 2004).

The absence of differences between sexes means that both females and males can explore novel environments, foods and objects, and that they can both adapt to novel conditions at colonisation fronts.

### Perspectives

In a study on the Iberian wall lizard, *Podarcis hispanica*, Rodriguez-Prieto et al. (2011) conducted a novel environment test and found individual differences in boldness. In repeated tests, they observed that there were consistent personalities in individuals, but also a habituation phenomenon. Individuals that were bolder habituated faster to the apparatus than shyer ones (Rodriguez-Prieto et al. 2011). Thus, habituation processes should also be studied in invasive species to see if the primary fear reactions of individuals who did not approach the object would be maintained in the long term or if experience enhances progressive colonization with a decrease of neophobia and emotional reactions. In another series of experiments, we observed that starlings rapidly habituate to the novel objects and can even learn to manipulate them in order to obtain food (Rodriguez 2010b). It is also important to understand what levels of boldness are adaptive and when boldness becomes costly: the bold Namibian rock agamas, *Agama planiceps* are reported to be more easily trapped than the shy ones (Carter et al. 2012,) and unreactive birds are hit by cars more often (Møller and Erritzøe 2017)

The presence of conspecifics can enhance or inhibit approaching and touching objects (Stöwe et al. 2006). Social aspects involved in neophobia need to be studied in order to better understand novelty-approaching dynamics in highly gregarious species like this one.

## Conclusion

Introductions of starlings in different countries involved groups of 60 to 100 individuals (Flux and Flux 1981, Feare 1984). The European starling, *Sturnus vulgaris* has successfully established self-sustaining populations in several of the regions where it has been introduced in North America, Australia, South Africa, New Zealand and Argentina (Pell and Tidemann 1997, Peris et al. 2005). We suggest that the diversity of reactions towards novelty, the presence of bold individuals in the groups, combined with social facilitation, enhanced the colonization processes allowing for the exploration of novel habitats, the consumption of novel foods and approaching novel objects (in particular in urban environments). The European starling did not invade habitats like forests and desert regions. We think that in these cases landscape structure is involved as starlings need a combination of trees to nest and open field areas to forage and escape from predators. More studies in the laboratory and in the field are required to understand which of the different existing profiles retard or accelerate invasion success.<colcnt=7>

Ethical Statement should be divided into the following sub-section:

- Funding
- Conflict of Interest
- Ethical approval
- Informed consent

## Ethical Statement

This study was supported by the Ministry of French Research by an Excellence Grant allocated for Doctor Alexandra Rodriguez (Bourse MRT) and by the laboratories of Ecology and Ethology Research INRA Scribe (ancient team of Ecology of Invasions) and CNRS UMR 6552 Ethos (Ethologie Animale et Humaine). The study takes into account ecology and socio-biology debates and postures around invasions biology. We used enough starlings to be able to conduct non parametrical tests and the maximum-minimum of birds required to fulfil the three Rs principle. This manuscript has been written, corrected and approved by its auhors.

## Acknowledgements

We thank Christine Aubry and Christophe Petton for taking care of the birds. We thank Dr. Adrian Craig who corrected the English language. The study was funded by a grant from INRA Scribe Ecology of Biological Invasions, CNRS and Rennes 1 University. Alexandra Rodriguez was supported by a PhD grant from the French Ministry of Research

